# Intense light unleashes male-male courtship behavior in wild-type *Drosophila*

**DOI:** 10.1101/2022.07.12.499756

**Authors:** Atsushi Ueda, Abigayle Berg, Tashmit Khan, Madeleine Ruzicka, Shuwen Li, Ellyn Cramer, Atulya Iyengar, Chun-Fang Wu

**Author notes:** Correspondence: Atsushi Ueda Chun-Fang Wu Atulya Iyengar.

## Abstract

*Drosophila* courtship studies have elucidated several principles of the neurogenetic organization of complex behavior. Through an integration across sensory modalities, males perform stereotypic patterns of chasing, courtship song production, and copulation attempts. Here we report a serendipitous finding that intense light not only enhances courtship toward female targets but also triggers unexpected courtship behaviors among male flies. Strikingly, in wild-type male-only chambers, we observed extreme behavioral manifestations, such as “chaining” and “wheeling”, resembling previously reported male-male courtship behaviors in *fruitless* mutants and in transformants with ectopic *mini-white*^*+*^ overexpression. This male-male courtship was greatly diminished in a variety of visual system mutants, including disrupted phototransduction (*norpA*), eliminated eye-color screening pigments (*white*), or deletion of the R7 photoreceptor cells (*sevenless*). However, light-induced courtship was unhampered in wing-cut flies, despite their inability to produce courtship song, a major acoustic signal during courtship. Unexpectedly the olfactory mutants *orco* and *sbl* displayed unrestrained male-male courtship. Particularly, *orco* males attained maximum courtship scores under either dim or intense light conditions. Together, our observations support the notion that the innate male courtship behavior is restrained by olfactory cues under normal conditions but can be unleashed by strong visual stimulation in *Drosophila*.

## Introduction

The male courtship behavior repertoire in *Drosophila melanogaster* emerges from a complex integration of external sensory information (1–3), prior experiences (4–6), and internal drive (7–9). A large collection of discrete visual (10–12), olfactory (13–15), gustatory (16–18) and mechanosensory (19) cues serve to direct courtship behavior towards receptive females. These signals in-turn trigger stereotypic sequences of chasing, licking, wing extension, courtship ‘song’ production and eventual mounting (20, 21). Genetic perturbations in this sensory-motor integration process can reveal contributions of the genetic-organizations and neural circuit-mechanisms underlying this complex behavioral program (22–24).

A striking phenotype arising from certain genetic manipulations is male-male courtship behaviors. For example, several *fruitless* mutants display remarkable ‘chains’ of courting males (25, 26). The *fruitless* gene product undergoes alternative splicing to produce female- and male-specific transcripts, *fru*^*F*^ and *fru*^*M*^, respectively (27, 28). In *fru*^*M*^-disrupted males, male-male courtship is often observed (27). Disruption of subsets of *fru*^*M*^-positive sensory neurons hampers reception of the male-specific repulsive cues, 7-tricosine and cis-vaccenyl acetate, leading to male-male courtship (29–33). Furthermore, it has been reported that optogenetic activation of *fru*^*M*^-expressing neural circuits induces male-male courtship (34). Male-male courtship has also been reported in studies using *painless* TrpA channel mutants (35), dopaminergic signaling mutants (36, 37), or transformants for forced expression of mini-white constructs (38, 39).

Here, we describe a serendipitous finding of male-male courtship behaviors in wild-type (WT) *Drosophila melanogaster* evoked by intense light. We observed the light intensity-dependent increase in chasing, wing extension, courtship song, and chaining in different strains of WT males, using LED-, incandescent-, and sun-light illumination. We examined the phenomena in flies with genetic and surgical manipulations to elucidate the key roles of the visual and olfactory systems in gating this male-male courtship behavior.

## Results

### Intense light riggers courtship behavior in male flies

A modified semi-automated Drosophila tracking system, IowaFLI Tracker (40) was employed to analyze activity in each circular arena containing eight flies (Figure 1). The tracking system consisted of a clear polyacrylic sheet containing four arenas which were housed in a light-shielded cylindrical chamber equipped with an inner circular strip of LED lighting (Figure 1A). A webcam mounted on top of the ceiling captured fly behaviors. In male-only arenas, we unexpectedly discovered a high frequency of courtship behaviors at intense light setting. Seconds after light on, we regularly observed chasing and attempting to court among male flies. Strikingly, we often saw chains of courting males (Figure 1B, Movie 1), reminiscent of phenotypes previously found in mini-white transgenic lines (38, 39) and *fruitless* mutants (25, 41). To quantify light-induced courtship, we developed a protocol where the eight flies in the arena were first subjected to 2-min illumination of relatively low intensity (0.4 klx, within the normal range of room lighting) followed by 2-min of high-intensity light (18 klx) and subsequently, another 2-min low-intensity light period (Figure 1D). For each successive 10-s interval, we manually scored the occurrence (0 or 1) of chasing, wing extensions and chaining in the arena. The total number of intervals during which the respective behaviors were observed in the arena was reported as the “total score” (ranging from 0 to 12 for the 2-min period, Figure 1D&E; see Methods). During the high-light period, the increase in total chasing score was more than 700%, ∼400% for wing extension, and greater than 800% for chasing (Figure 1E).

**Fig. 1.**
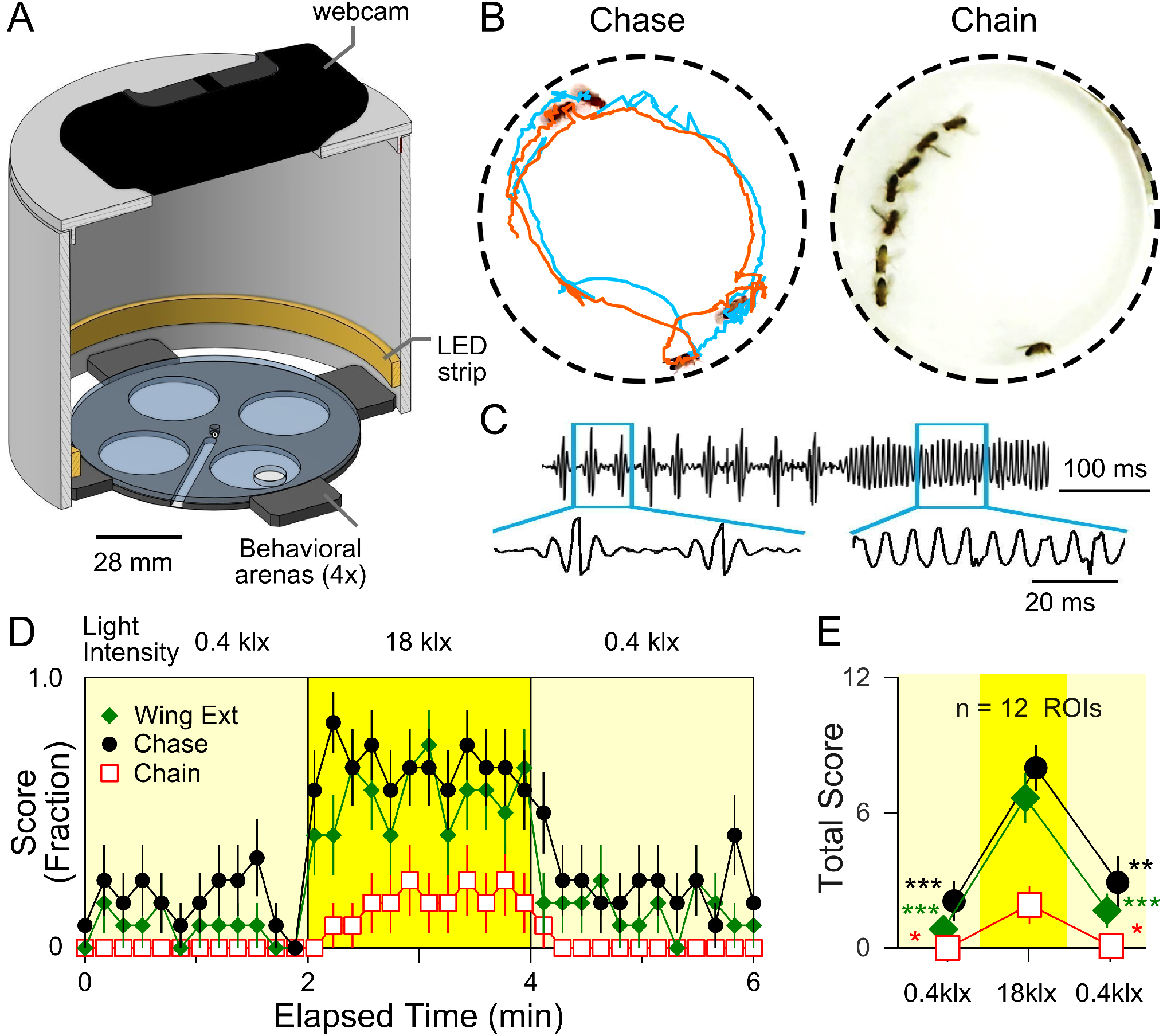
Light-evoked male-male courtship behavior in *Drosophila*. (A) Schematic drawing of the behavioral arena, LED strip lighting and webcam configuration. Eight males were loaded into each of the four circular behavioral arenas (regions of interest, ROIs). An LED strip surrounding the arenas illuminated fly behavior with minimal shadowing, and a webcam above recorded activity. (B) Intense light triggers male-male courtship behaviors including chase (left, orange track chasing cyan track), as well as chain and wing extensions (right). (C) In a modified arena equipped with a microphone (see Methods), intense light triggered pulse and sine songs among male flies, resembling qualitatively songs recorded during male-female courtship (Fig. S1). (D) Quantification of light-induced male-male courtship behaviors. Eight males were subjected to a 2-min low-light intensity period, followed by a 2-min high-intensity light, and a 2-min low-light recovery period. For each successive time bin (10 s), the presence or absence of respective behaviors was scored (1 or 0). Data points presented show the average score pooling from 12 ROIs for the various behaviors (see Methods). Error bars indicate SEM. (E) Total scores in D are summed over 2-min periods and shown with SEM and # ROIs. One-way ANOVA, Bonferroni-corrected t-test post hoc was applied for comparisons between 1st low-light and high-light, as well as between high-light and the 2nd low-light. *p < 0.05, **p < 0.01, ***p < 0.001.

In a modified arena equipped with a microphone, we confirmed the light-induced wing extension behavior was related to song production among male flies (Figure 1C & S1). We found that like male-female courtship songs, male-male songs consisted of pulse and sine songs. The frequency and duration of sine songs as well as the inter-pulse intervals and number of pulses per bout of the pulse song were analyzed, showing overlapping ranges of these parameters between male-male and male-female songs (Figure S1B).

### Generality and specificity of light-induced courtship behaviors

To determine whether light-induced male-male courtship is a specific phenomenon restricted to our WT *Canton-S* (*CS*) strain used in most of the experiments, we tested another WT strain of *melanogaster, Berlin*, and found the same male-male courtship behaviors as shown in Figure 2A. We observed similar high-light-induced increases in wing extension and chasing behavior as in *CS* flies, with occasional chaining observed as well (Figure 2A). We also observed mixed-sex arenas (4 male and 4 female flies) to examine how high-intensity illumination affects male-female interactions, to determine whether the intense light-triggered courtship behavior was restricted towards other males only. As shown in Figure 2B, in these mixed-sex arenas, high-intensity lighting also similarly increased male-female courtship behaviors (∼300%). In a subset of mixed sex arenas, we determined the kinetics of intense-light effect on male-female courtship scores and found temporal properties similar to male-only arenas (Figure S2). Together, these findings suggest intense light intensifies courtship drive of male flies toward both male and female flies, rather than switches their sex preference from females to males.

**Fig. 2.**
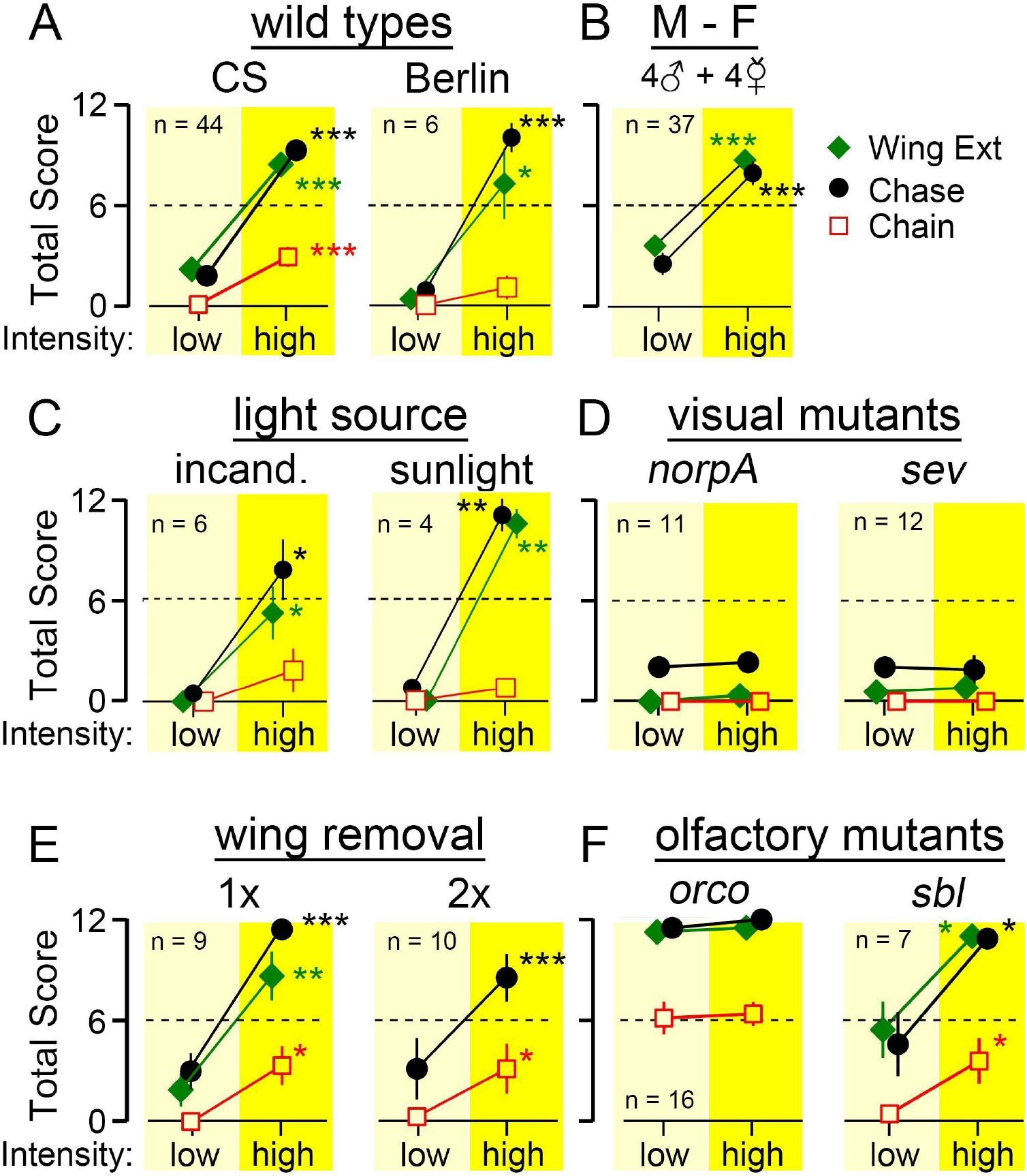
Genetic and environmental factors affecting light-evoked male-male courtship. Flies were subjected to a 2-min period of low light intensity followed by 2-min of high light intensity (0.4 klx and 18 klx respectively, except for panel C with different light sources). (A) Light-evoked male-male courtship in the CS and Berlin WT *Drosophila* strains. (B) Courtship behavior in mixed sex arenas with 4 males and 4 females. (C) Male-male courtship in arenas under incandescent light (left) or in direct sunlight (right). For incandescent illumination, light was delivered by two light pipes from an USHIO EKZ 10.8 V, 30 W bulb (‘low’ intensity: 0.4 klx, ‘high’ intensity 25 klx). For sunlight experiments, the behavioral arena was placed in a shaded area for the ‘low’ period (0.4 klx) and moved to direct sunlight (90 klx) for ‘high’ period. Experiments were done in the Biology Building courtyard on the University of Iowa campus on 19/05/2022 at 15:00 CDT (temp 29 °C). (D) Lack of light-triggered courtship behaviors in visual mutants (norpA and sev). (E) Wing-cut flies and (F) Olfactory mutants (orco and sbl). Note the light-induced increase in courtship behavior is absent in visual mutants, but unimpaired in flies with one (1x) or both (2x) wings removed. Olfactory mutants show greatly enhanced courtship activity, with orco attaining nearly maximum scores even at low-light intensity. Data are shown as mean +/- SEM. * p < 0.05, ** p < 0.01, *** p < 0.001, paired t-test (high vs. low). For each panel, the sample size (ROIs) is indicated.

### Roles of sensory cues in light-triggered male-male courtship

We characterized the efficacy of different lighting conditions for triggering the male-male phenomenon. By adjusting the arena LED light intensity, we found that the relationship between light intensity and incidence of wing extensions, chasing and chaining behaviors was monotonic and that the proportionally steepest increases occurred between 2.0 and 6.0 klx (Figure S3). We then asked whether the LED light source we used was specifically required, or some other sources could also evoke the effect. As shown in Figure 2C, intense incandescent light (25 klx) also elicited courtship behavior. We further asked if natural light (e.g., sunlight on a spring day) could induce male-male courtship activity. Remarkably, after moving the arena from a shaded location (∼0.4 klx) to a location under direct bright sunlight (∼90 klx), we observed robust courtship activity (Figure 2C). These results indicate that a wide variety of sufficiently intense, broad-spectrum visible light sources (see Figure S4 for spectral measurements of different light sources) can effectively induce male-male courtship behavior.

We introduced experimental perturbations to the various sensory systems for indications of their roles in the light-induced male-male courtship. First, we examined males with a disrupted visual system. The gene *norpA* encodes the enzyme phospholipase C required for phototransduction, and several mutations eliminate photoreceptor potential, rendering mutant flies completely blind (42). In blind *norpA*^*P12*^ mutants, we found intense light did not induce courtship activity (Figure 2D, Movie 1).

We further took advantage of anatomical mutations affecting a particular category of photoreceptors. The *Drosophila* compound eye is composed of about 800 facets, each cylindrical rhabdomere below consisting of 6 outer photoreceptors (R1 - R6, green light sensitive) and two central photoreceptors (R7 and R8, UV and blue light, respectively). In *sevenless* (*sev*) mutants, the high-acuity channels composed of central R7 and R8 are functionally impaired because R7 is absent, which also disrupts the light guide for R8 underneath, while the high sensitivity channel, consisting of R1-R6, is intact (43). The results showed that light-evoked male-male courtship was absent in *sev* mutants (Figure 2D), implying a critical role for the R7/R8 system in light-triggered male-male courtship. Visual processing depends on integration of photoreceptor responses from individual rhabdomeres. Because of the screening pigments, each rhabdomere acts as an independent light guide for input from a narrow angular visual field, in isolation of its neighbors (44). Mutations of the *white* (*w*) gene are devoid of the screening pigments and thus degrade visual image processing, in spite of increased sensitivity to light (45). In *w* mutant alleles, including *w*^*1118*^ and *w*^*G*^, despite their active locomotion in the arena, the male-male courtship behavior was missing (Figure S5). Conceivably, visual acuity is crucial to the locomotion control during courtship. It is documented that *white* mutant males show lower performance in male-female courtship (46, 47). Further, additional eye color mutants have been reported to be defective in courtship behavior (48). We have examined *cinnabar* (cn) and *cinnabar brown* (*cn bw*). The *cn* flies lack the screening pigment ommochrome (49) and show bright orange eye color; *bw* lack pteridine (49) and *cn bw* flies are white-eyed. Our initial observations suggest that both *cn* and *cn bw* mutants showed greatly decreased intense-light-induced male-male courtship, comparable to the *w* alleles described above. These observations indicate the central role of visual function in the light-evoked male-male courtship.

In contrast to the visual system, manipulations of mechanosensory or auditory cues revealed drastically different outcomes. In groups of males, courtship behaviors are known to be triggered by playback of courtship song (19). However, we found that in wing-cut male flies, which had disrupted wing mechanosensory inputs and were unable to produce courtship song, intense light nevertheless reliably triggered chasing and chaining (Figure 2E). Notably, flies with one wing or both wings cut displayed similar chasing and chaining scores and flies with only a single wing could attain a frequency of wing-extension comparable to WT. This result demonstrates that the auditory cues generated by wing beats and the mechanosensory receptors along the wing blades are not required in the light-evoked male-male courtship.

To determine the role of olfactory systems, we examined two mutants *smellblind* (*sbl*), a *para* Na^+^ channel allele with disrupted odor-evoked behaviors (50) and *odorant receptor co-receptor* (*orco*), encoding a requisite olfactory receptor subunit (51). Surprisingly, at low light intensities, both mutants displayed strikingly elevated courtship behaviors compared to WT males (Figure 2F). Particularly, recurrent male-male courtship in *orco* flies was so prevalent (Movie 1), frequently approaching the maximum score of 12, that a further increase in courtship was not apparent during the intense light periods. However, in *sbl* males, which showed milder increases in low-light scores, intense-light could further promote courtship activities to attain the maximum score (Figure 2F), prompting a further investigation into interactions between intense light and olfactory processing in gating male courtship behaviors.

Apparently, olfactory processing plays a similar role in both male-male and male-female courtship behaviors (Figure S6). We monitored the intense-light effect on male-female courtship in *sbl* and *orco* mutant flies. Consistent with male-male courtship, *sbl* male-female courtship behavior was further increased by intense light, while a less obvious effect on *orco* was observed because their chasing and wing extension scores were already near saturation at the low light intensity.

### Spatio-temporal properties of male-male courtship behaviors

In addition to courtship event statistics shown in Figure 2, we also monitored continuous activity patterns among the male flies, using IowaFLI Tracker (40) for an automated analysis of kinematic properties as well as social interactions among individually identified flies. As Figure 3A shows, WT flies, upon high-intensity illumination, displayed a drastic increase in activity levels (panel i, locomotion tracks, with individual flies color-coded) as well as a greatly enhanced level of social interactions (panels ii, interactograms, with colored time segments registering other individuals active within the 3.75 mm vicinity. An interaction proximity criterion of 3.75 mm between centroids was adopted because it optimally captures chasing events in WT flies.). In contrast, *norpA* flies, upon high-intensity illumination, displayed neither a drastic increase in activity levels (panel i) nor a greatly enhanced level of social interactions (panel ii). No male-male chasing or wing extension events were evoked by intense light (Figure 2D). Interestingly, *orco* flies exhibited a contrasting case for reduced response to intense light. Chasing or chaining events already occurred at low-light intensities, indistinguishable from behavioral activities under intense light, as indicated by the locomotion tracks (panel i) and interactograms (panel ii).

**Fig. 3.**
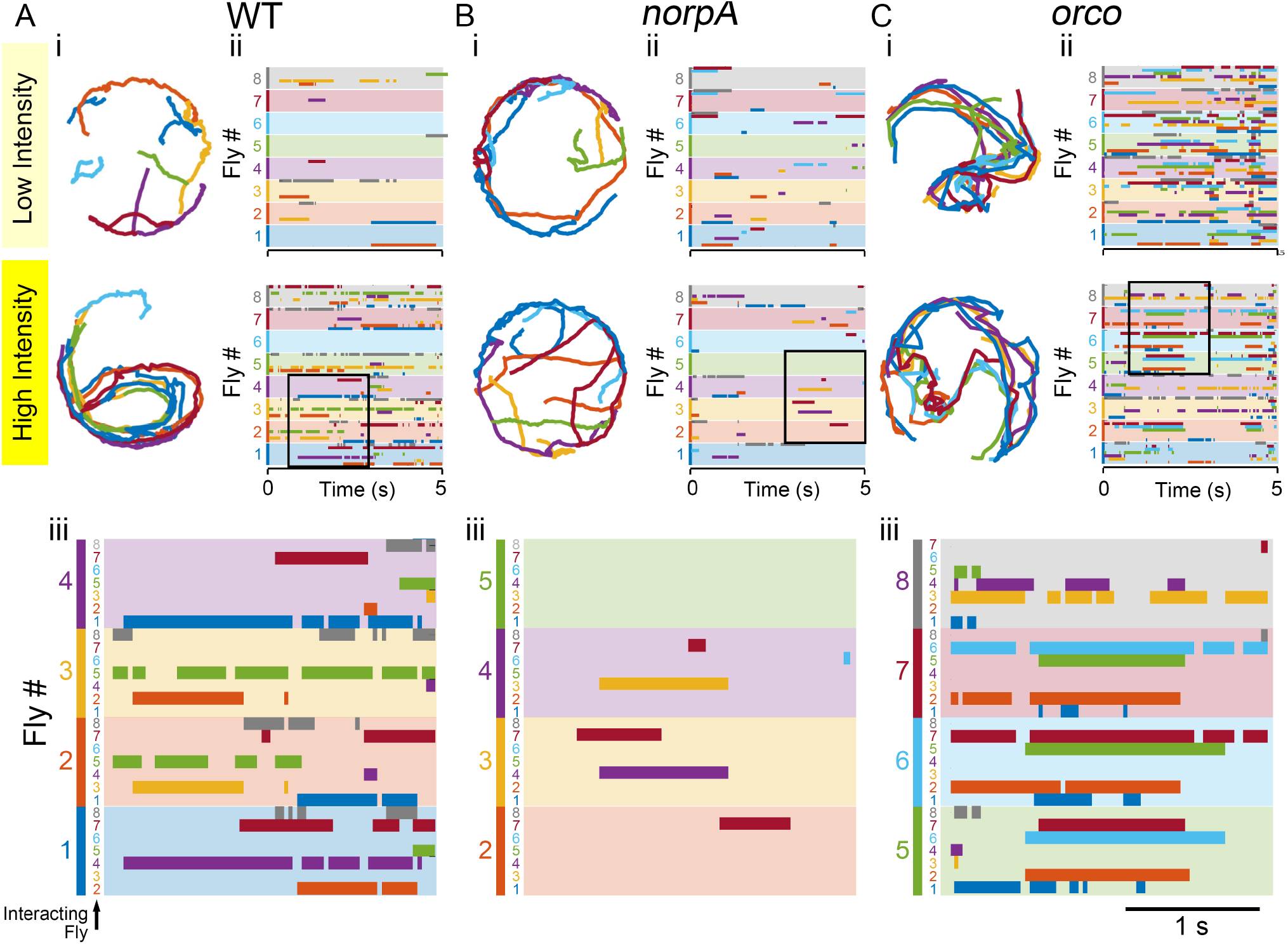
Fly tracks and interactogram of light-induced male-male courtship. Representative behavioral trajectories (i; left panels for each genotype; 5-s sample) and corresponding interactograms (ii; right panels) of WT, *norpA*, and *orco* males under low and high light. Note that flies are chasing others under high light in WT, and under both low and high light in *orco* mutants. However, *norpA* flies displayed no increase in levels of locomotion or social interactions. To monitor fly activity patterns, the IowaFLI Tracker (40) was used for automated analysis of kinematic properties of individual flies as well as social interactions among them. An interaction proximity criterion of 3.75 mm, optimal for capturing chasing events, was adopted. For each genotype, individual flies (#1-8) were color-coded to depict their continuous locomotion tracks (i), and to register events of its interactions with others in the interactogram (each color-coded, falling within the 3.75-mm mutual distance) along the 5-s duration (ii). In the horizontal shaded bands of 8 different colors, interaction of the fly of the designated band color with all other individuals were registered with darker color-coded line segments to indicate the time intervals when their mutual distance < 3.75 mm. (iii) A selected area (boxed) in the interactogram is enlarged to better illustrate the reciprocal relationship among the interacting individuals identifiable with assigned colors.

In Figure 3, a selected area (boxed) in the interactogram panel for each genotype is enlarged to illustrate interaction records for individually identified flies (panel iii). Each fly in the arena was assigned a number and a color code. In each color-shaded strip for an identified fly, the other flies in the vicinity of this individual are recorded as bar segments of darker colors. Thus, the reciprocal relationship among the color-coded individuals can be traced for each time point in the original video record (15 frames/s). The interactogram provides the raw source data for time-stamped proximity information for the involved individuals. Following visual inspection, particular categories of courtship interactions (wing extension, chasing, and chaining) could be related back to the video recording and the locomotion tracks to obtain a more comprehensive description of the event.

IowaFli Tracker also provides statistics for kinematics parameters, such as distance travelled, speed, and % time moving (Table S1). From the table, *norpA* is clearly hyperactive (with high values of speed, distance travelled, and % time moving). However, most of the recorded *norpA* interactions, defined by the 3.75-mm proximity criterion, were not courtship-related and occurred most often along the arena periphery (Figure 3Bi). In contrast, many interactions indicated in the interactograms for WT and *orco* could be related to compact bundles of locomotion tracks traversing across the arena for the same time segments (Figure 3Ai and ii, Ci and ii).

## Discussion

### The role of visual system function in male-male courtship

Our findings add a striking case of male-male courtship behavior in WT flies, in contrast to previous reports from a number of genetic variants, including *fru* (25, 41), *painless* (35), an unidentified mutation (on Chr. 3L) reported by Sharma (52), as well as several mini-*w*^*+*^ transformants (38, 39). A strong visual contribution to such behaviors was suggested by Sharma’s observation that blue light inhibits male-male courtship (52). Similarly, blue and UV lights have been reported to suppress male-female courtship (6). In a preliminary follow-on experiment, we subjected *orco*^*1*^ and *orco*^*2*^ males to intense blue light and found that male-male courtship was severely suppressed (Table S2). It is important to note that the two *orco* alleles employed in this study carry the mini-w^+^ construct (in place of the *orco* gene) in a w^-^ background (See Methods for the construct). Their eye color appeared normal, indicating strong mini-w^+^ expression. Therefore, the extreme phenotypes of *w;;orco*^*1*^ and *w;;orco*^*2*^ in this report may have had contributions from mini-w^+^ expression.

Our preliminary analysis of *orco*^*1*^ and *orco*^*2*^ heterozygotes (*orco*/+) generated from crossing *w/Y; orco* males to *w*^*1118*^ females displayed a correlation between eye-color density and male-male courtship activity. We observed darker eye color in *w;;orco*^*1*^*/+* than *w;;orco*^*2*^*/+*, indicating differences in mini-w^+^ expression, and accordingly a stronger male-male courtship behaviors in *w;;orco*^*1*^*/+* (Table S3). In terms of male-male courtship scores, we observed in the following sequence as shown in Table S3:

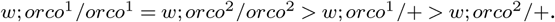

Furthermore, *norpA*^*P12*^*/Y;;orco*^*1*^*/+* and *sev*^*LY3*^*/Y;;orco*^*1*^*/+* males have been obtained and the behavior test suggests total suppression of *orco*^*1*^*/+* phenotype, behaving like *norpA*^*P12*^ and *sev*^*LY3*^ males (Table S3). Taken together, our results confirm a critical role of visual system function in male-male courtship.

### The effective light intensity range and underlying candidate neural circuits

The above findings demonstrate that light plays a major role in courtship behavior in *Drosophila melanogaster*. Besides facilitating visually guided motor behaviors, intense light serves as a potent modulator promoting male sex drive as indicated by increased courtship behavior towards males as well as females. Notably, the light intensity range required to effectively trigger male-male courtship (between 2 and 6 klx, Figure S3) is several fold greater than common light sources encountered in the lab, including room lights or incubator lighting (0.5 klx, measurements in our lab). However, this light intensity is within the range that flies encounter in the wild, as direct sunlight can be above 100 klx (53). Importantly, our findings with both incandescent light and sunlight (Figure 2C) indicate a variety of sufficiently intense broad-band light sources (Figure S4) can induce male-male courtship. It is tempting to speculate based on recent advances in identification and manipulation of neuronal components of various circuits involved in male courtship behavior. Our analysis suggests an interplay of visual and olfactory processing. Indeed, in a model previously proposed by Wang et al. (35), male flies have an intrinsic drive to court that is normally restrained by olfactory processing of certain signals from other males (e.g., cis-vaccenyl acetate (22)). Flies deficient in olfactory processing such as *orco* and *sbl* would lack these restraints, consistent with the increased male-male courtship shown in Figures 2 and 3. Conceivably, intense light could serve to unleash the intrinsic courtship drive either by supercharging the courtship motor program to overcome olfactory inhibition or by directly suppressing the inhibitory olfactory processing in WT flies. Within the broad-band illumination used in this study, the shorter wavelength UV and blue light preferentially activate R7 and R8, respectively. UV and blue light are known to suppress male-female courtship (6) and blue light was found to inhibit male-male courtship in the mutant flies reported by Sharma (52). Our analysis of *norpA* and *sev* mutants implies that R7 and/or R8 processing plays an important role in light-induced courtship (Figure 2D). Photoreceptors R7 and R8 directly target several classes of medulla neurons, including Dm8 amacrine-like cells, and Tm5 and Tm9 cells which project to the lobula plate (54). It is possible that these medulla neurons directly or indirectly influence excitability of fru-expressing P1 neurons in the lobula plate, which exert strong modulation on the activity of LC10a circuit that has been shown to be essential for controlling courtship chasing performance (3, 55). Future analysis will elucidate the action and modulation of the neural circuit underlying light-evoked male-male courtship. Our findings provide a new context of courtship behavior and can add constraints for data interpretation in the studies of neural circuits responsible for the repertoires in male courtship behaviors.

## Methods

### *Drosophila* Stocks and Husbandry

All flies were reared at room temperature (23 °C) under a 12:12 light-dark cycle in vials containing cornmeal media(56, 57). All flies studied were age-controlled as specified. Throughout, WT strain used was *Canton-S* (except for Figure 2A, where WT-*Berlin* was used, gift from M. Heisenberg (44)). Other mutant lines studied included: *smellblind* (*sbl*^*1*^, *sbl*^*2*^), EMS-induced mutant alleles of the voltage-gated sodium channel gene *paralytic* (gift from J. Carlson(50)); *orco*^*1*^ and *orco*^*1*^ (*w*;; TIw+mW*.*hs=TIorco*^*1*^ and *w*;; TIw+mW*.*hs=TIorco*^*2*^; Bloomington 23129 and 23130), two insertion alleles of the gene *odorant receptor co-receptor* (*orco*); *norpA*^*P12*^, an EMS-induced allele of *no receptor potential A*, encoding a phospholipase C required for visual transduction (gift from W.L. Pak(42)); *sevenless*^*LY3*^ (*sev*^*LY3*^) lacking R7 photoreceptors (gift from W.L. Pak(58)); *cn*^*1*^ (Bloomington 263); and two alleles of *white, w*^*1118*^ (Bloomington 6326) and *w*^*G*^ (gift from M.L. Joiner).

### Courtship Behavioral Recording

Fly courtship behavior was observed in a custom-built chamber which consisted of an arena piece, and arena ceiling cover, and a light-shielded cylindrical chamber (Figure 1A). The arena piece consisted of a plexiglass sheet (2 mm thick) with four milled circular chambers referred to as “Regions of Interest” or ROIs, (26 mm diameter) seated on filter paper (Whatman #1) forming 2 mm-high behavioral arenas (4 ROIs). The arena ceiling cover was a plexiglass sheet of the same specification. The light-shielded chamber was constructed from a 4” PVC pipe coupler (NIBCO #4801) with a custom milled PVC lid. A 21 LED strip (STN-A30K80-A3A-10B5M-12V, SuperBrightLEDs.com; Figure S4) was glued along the inner rim next to the base of the light chamber. LEDs were powered by a custom-built current-controlled power supply (Figure S5). For Figure 2C experiments, incandescent light was delivered via fiber optic light pipes from an USHIO EKZ 10.8 V, 30 W bulb. Spectral characteristics of the above light sources were determined across a range of power supply currents using a Flame Miniature Spectrometer (Ocean Optics, Orlando, FL). The spectral power across a range of wavelengths (λ =189 nm to 884 nm) was measured following manufacturer’s instructions. In Figure S4, spectral data were normalized to the peak emission detected (i.e. E(λ) / E(peak) for the wavelength λ). In addition, the spectral characteristics of sunlight on a sunny spring day (2PM, April 10, 2023, in Tuscaloosa, AL, USA) was measured (Figure S4).

Fly behaviors were captured by a Logitech c920 webcam connected to a PC. For microphone recordings (Figure 1B), a modified arena of a larger dimension was constructed with a hole drilled in the side of the arena. A high-gain microphone (FC-23629-P16, Knowles Electronics, Itasca, IL, USA; see also(59)) was placed in the opening to pick up fly sounds. Light intensity measurements (Figure S2) were performed using a digital light meter (DLM1, Universal Enterprise Inc. Portland, OR, USA) following manufacturer’s instructions. Behavioral recordings were performed by aspirating 8 flies into each arena (without anesthesia). Following a brief acclimatization period (< 5 min), flies were recorded for 2 min under ‘low-light’ conditions, immediately followed by 2-min of ‘high-light’ conditions. In some experiments (Figure 1C-D), we included a 2-min ‘low-light’ recovery period.

### Quantification and Statistical Analysis

Behavioral scoring was carried out in groups of 8 flies confined in the arena. (For male-female courtship, 4 males and 4 females were used). Criteria for chasing and wing extension behaviors were based on Hall(20), and we define chaining behavior as the formation of a mobile chain of 4 or more flies (cf. Zhang & Odenwald(38) and Hing & Carlson(39)). Scoring was performed by a pair of operators (an observer and a recorder), either on-site during live video recording or by viewing play-back of videos. Recording and scoring were performed on pairs of ROIs, one for experimental and one for control (or comparison) flies. Each successive 10-second interval in the 2-min observation periods was divided into two 5-second scoring windows and experimental and control ROIs were scored alternately (i.e., 5 s for one ROI followed by 5 s for the other). the presence or absence of each of the courtship behaviors in an ROI was recorded in the 5-second window. The total number of 5-s windows (out of 12 possible) in which the respective behaviors were observed, reported as the “total score,” ranging from 0-12. In addition, automated analysis of walking kinematics was performed offline with IowaFLI Tracker as described in Iyengar et al.(40).

All statistical analyses were performed in MATLAB r2021b. Unless otherwise indicated, all statistically significance values are reported from paired t-tests comparing the ‘low’ vs ‘high’ light conditions. For comparisons across multiple groups (Figures 1E, S2D), a one-way ANOVA analysis was performed as a preliminary step. Table S4 indicates the complete list of statistical analyses and results in this study.

## ACKNOWLEDGEMENTS

We thank the participants of the 2019, 2020, 2022 Neurogenetics Lab course at University of Iowa and the course TA, Quinn Christensen for their assistance in data collection. We also thank Abigail Hawken, Bryan Joos, and Crystal Kim for additional data collection. We are grateful for the assistance of Jeremy Richardson and the Biology Engineering Shop in fabricating behavioral arenas. We thank Kyle Edwards and Eddy Lontchi of the Dixon Lab at University of Alabama for their assistance in measuring light source emission spectra. This work was supported by an NIH R01 AG 051513 and NS 1111122 to CFW and an Iowa Neuroscience Institute Fellowship to AI.

## Ethics

N/A

## Data, code and materials

An updated version of IowaFLI Tracker (version 3.1) and the original data sets for figure and table construction can be found on github (*https://github.com/IyengarAtulya/IowaFLItracker*).

## Contributions

A.U.: conceptualization, data curation, formal analysis, investigation, methodology, supervision, visualization, writing-review & editing; A.B.: formal analysis, investigation, supervision; T.K.: conceptualization, investigation, visualization; S.L.: investigation; M.R.: formal analysis, investigation, visualization; E.C.: investigation; C.F.W.: conceptualization, funding acquisition, investigation, methodology, project administration, visualization, writing - review & editing; A.I. conceptualization, data curation, formal analysis, investigation, methodology, resources, software, supervision, validation, visualization, writing – original draft.

## Bibliography

1. Ralph J Greenspan and Jean-François Ferveur. Courtship in drosophila. Annual review of genetics, 34(1):205–232, 2000.

2. Jean-François Ferveur. Cuticular hydrocarbons: their evolution and roles in drosophila pheromonal communication. Behavior genetics, 35:279–295, 2005.

3. E Josephine Clowney, Shinya Iguchi, Jennifer J Bussell, Elias Scheer, and Vanessa Ruta. Multimodal chemosensory circuits controlling male courtship in drosophila. Neuron, 87(5): 1036–1049, 2015.

4. William G Quinn and Ralph J Greenspan. Learning and courtship in drosophila: two stories with mutants. Annual review of neuroscience, 7(1):67–93, 1984.

5. Leslie C Griffith and Aki Ejima. Courtship learning in drosophila melanogaster: diverse plasticity of a reproductive behavior. Learning & Memory, 16(12):743–750, 2009.

6. Takaomi Sakai, Kunio Isono, Masatoshi Tomaru, Akishi Fukatami, and Yuzuru Oguma. Light wavelength dependency of mating activity in the drosophila melanogaster species subgroup. Genes & genetic systems, 77(3):187–195, 2002.

7. Stephen X Zhang, Dragana Rogulja, and Michael A Crickmore. Recurrent circuitry sustains drosophila courtship drive while priming itself for satiety. Current Biology, 29(19):3216–3228, 2019.

8. Weiwei Liu, Anindya Ganguly, Jia Huang, Yijin Wang, Jinfei D Ni, Adishthi S Gurav, Morris A Aguilar, and Craig Montell. Neuropeptide f regulates courtship in drosophila through a male-specific neuronal circuit. Elife, 8:e49574, 2019.

9. Selim Terhzaz, Philippe Rosay, Stephen F Goodwin, and Jan A Veenstra. The neuropeptide sifamide modulates sexual behavior in drosophila. Biochemical and biophysical research communications, 352(2):305–310, 2007.

10. Sweta Agrawal and Michael H Dickinson. The effects of target contrast on drosophila courtship. Journal of Experimental Biology, 222(16):jeb203414, 2019.

11. Robert Cook. The extent of visual control in the courtship tracking of d. melanogaster. Biological Cybernetics, 37(1):41–51, 1980.

12. Ken-ichi Kimura, Chiaki Sato, Kana Yamamoto, and Daisuke Yamamoto. From the back or front: the courtship position is a matter of smell and sight in drosophila melanogaster males. Journal of neurogenetics, 29(1):18–22, 2015.

13. Sandeep Robert Datta, Maria Luisa Vasconcelos, Vanessa Ruta, Sean Luo, Allan Wong, Ebru Demir, Jorge Flores, Karen Balonze, Barry J Dickson, and Richard Axel. The drosophila pheromone cva activates a sexually dimorphic neural circuit. Nature, 452(7186): 473–477, 2008.

14. Donald A Gailey, Robert C Lacaillade, and Jeffrey C Hall. Chemosensory elements of courtship in normal and mutant, olfaction-deficient drosophila melanogaster. Behavior genetics, 16:375–405, 1986.

15. Yael Grosjean, Raphael Rytz, Jean-Pierre Farine, Liliane Abuin, Jérôme Cortot, Gregory SXE Jefferis, and Richard Benton. An olfactory receptor for food-derived odours promotes male courtship in drosophila. Nature, 478(7368):236–240, 2011.

16. Hubert Amrein. Pheromone perception and behavior in drosophila. Current opinion in neurobiology, 14(4):435–442, 2004.

17. Jean-Francois Ferveur and Gilles Sureau. Simultaneous influence on male courtship of stimulatory and inhibitory pheromones produced by live sex-mosaic drosophila melanogaster. Proceedings of the Royal Society of London. Series B: Biological Sciences, 263(1373):967–973, 1996.

18. Claudio W Pikielny. Sexy deg/enac channels involved in gustatory detection of fruit fly pheromones. Science signaling, 5(249):pe48–pe48, 2012.

19. Eran Tauber and Daniel F Eberl. Song production in auditory mutants of drosophila: the role of sensory feedback. Journal of Comparative Physiology A, 187(5):341–348, 2001.

20. Jeffrey C Hall. The mating of a fly. Science, 264(5166):1702–1714, 1994.

21. Herman T Spieth. Courtship behavior in drosophila. Annual review of entomology, 19(1): 385–405, 1974.

22. Daisuke Yamamoto and Masayuki Koganezawa. Genes and circuits of courtship behaviour in drosophila males. Nature Reviews Neuroscience, 14(10):681–692, 2013.

23. Hania J Pavlou and Stephen F Goodwin. Courtship behavior in drosophila melanogaster: towards a ‘courtship connectome’. Current opinion in neurobiology, 23(1):76–83, 2013.

24. Barry J Dickson. Wired for sex: the neurobiology of drosophila mating decisions. Science, 322(5903):904–909, 2008.

25. Jeffrey C Hall. Courtship among males due to a male-sterile mutation in drosophila melanogaster. Behavior genetics, 8(2):125–141, 1978.

26. Adriana Villella, Donald A Gailey, Barbra Berwald, Saiyou Ohshima, Phillip T Barnes, and Jeffrey C Hall. Extended reproductive roles of the fruitless gene in drosophila melanogaster revealed by behavioral analysis of new fru mutants. Genetics, 147(3):1107–1130, 1997.

27. Ebru Demir and Barry J Dickson. fruitless splicing specifies male courtship behavior in drosophila. Cell, 121(5):785–794, 2005.

28. Lisa C Ryner, Stephen F Goodwin, Diego H Castrillon, Anuranjan Anand, Adriana Villella, Bruce S Baker, Jeffrey C Hall, Barbara J Taylor, and Steven A Wasserman. Control of male sexual behavior and sexual orientation in drosophila by the fruitless gene. Cell, 87(6): 1079–1089, 1996.

29. Tetsuya Miyamoto and Hubert Amrein. Suppression of male courtship by a drosophila pheromone receptor. Nature neuroscience, 11(8):874–876, 2008.

30. Seok Jun Moon, Youngseok Lee, Yuchen Jiao, and Craig Montell. A drosophila gustatory receptor essential for aversive taste and inhibiting male-to-male courtship. Current Biology, 19(19):1623–1627, 2009.

31. Beika Lu, Angela LaMora, Yishan Sun, Michael J Welsh, and Yehuda Ben-Shahar. ppk23-dependent chemosensory functions contribute to courtship behavior in drosophila melanogaster. PLoS genetics, 8(3):e1002587, 2012.

32. Robert Thistle, Peter Cameron, Azeen Ghorayshi, Lisa Dennison, and Kristin Scott. Contact chemoreceptors mediate male-male repulsion and male-female attraction during drosophila courtship. Cell, 149(5):1140–1151, 2012.

33. Shiu-Ling Chen, Bo-Ting Liu, Wang-Pao Lee, Sin-Bo Liao, Yao-Bang Deng, Chia-Lin Wu, Shuk-Man Ho, Bing-Xian Shen, Guan-Hock Khoo, Wei-Chiang Shiu, et al. Wake-mediated modulation of cva perception via a hierarchical neuro-endocrine axis in drosophila male-male courtship behaviour. Nature Communications, 13(1):2518, 2022.

34. Ryoya Tanaka, Tomohiro Higuchi, Soh Kohatsu, Kosei Sato, and Daisuke Yamamoto. Optogenetic activation of the fruitless-labeled circuitry in drosophila subobscura males induces mating motor acts. Journal of Neuroscience, 37(48):11662–11674, 2017.

35. Kaiyu Wang, Yanmeng Guo, Fei Wang, and Zuoren Wang. Drosophila trpa channel painless inhibits male–male courtship behavior through modulating olfactory sensation. PLoS One, 6(11):e25890, 2011.

36. Tong Liu, Laurence Dartevelle, Chunyan Yuan, Hongping Wei, Ying Wang, Jean-François Ferveur, and Aike Guo. Increased dopamine level enhances male–male courtship in drosophila. Journal of Neuroscience, 28(21):5539–5546, 2008.

37. Bin Chen, He Liu, Jing Ren, and Aike Guo. Mutation of drosophila dopamine receptor dopr leads to male–male courtship behavior. Biochemical and biophysical research communications, 423(3):557–563, 2012.

38. Shang-Ding Zhang and Ward F Odenwald. Misexpression of the white (w) gene triggers male-male courtship in drosophila. Proceedings of the National Academy of Sciences, 92 (12):5525–5529, 1995.

39. Audrey Liang Yin Hing and John R Carlson. Male-male courtship behavior induced by ectopic expression of the drosophila white gene: Role of sensory function and age. Journal of neurobiology, 30(4):454–464, 1996.

40. Atulya Iyengar, Jordan Imoehl, Atsushi Ueda, Jeffery Nirschl, and Chun-Fang Wu. Automated quantification of locomotion, social interaction, and mate preference in drosophila mutants. Journal of neurogenetics, 26(3-4):306–316, 2012.

41. Hiroki Ito, Kazuko Fujitani, KAZUE Usui, Keiko Shimizu-Nishikawa, Shoji Tanaka, and Daisuke Yamamoto. Sexual orientation in drosophila is altered by the satori mutation in the sex-determination gene fruitless that encodes a zinc finger protein with a btb domain. Proceedings of the National Academy of Sciences, 93(18):9687–9692, 1996.

42. Brian Thomas Bloomquist, RD Shortridge, S Schneuwly, M Perdew, C Montell, H Steller, G Rubin, and WL Pak. Isolation of a putative phospholipase c gene of drosophila, norpa, and its role in phototransduction. Cell, 54(5):723–733, 1988.

43. Satoko Yamaguchi, Reinhard Wolf, Claude Desplan, and Martin Heisenberg. Motion vision is independent of color in drosophila. Proceedings of the National Academy of Sciences, 105(12):4910–4915, 2008.

44. Martin Heisenberg and Reinhard Wolf. Vision in Drosophila: genetics of microbehavior, volume 12. Springer-Verlag, 2013.

45. Baruch Minke, Chun Fang Wu, and WL Pak. Isolation of light-induced response of the central retinula cells from the electroretinogram of drosophila. Journal of comparative physiology, 98:345–355, 1975.

46. Sheldon C Reed and Elizabeth W Reed. Natural selection in laboratory populations of drosophila. ii. competition between a white-eye gene and its wild type allele. Evolution, 4 (1):34–42, 1950.

47. Dimitrije Krstic, Werner Boll, and Markus Noll. Influence of the white locus on the courtship behavior of drosophila males. PloS one, 8(10):e77904, 2013.

48. Kevin Connolly, Barrie Burnet, and David Sewell. Selective mating and eye pigmentation: an analysis of the visual component in the courtship behavior of drosophila melanogaster. Evolution, pages 548–559, 1969.

49. JP Phillips and HS Forrest. Ommochromes and pteridines. Genetics and biology of Drosophila, 1980.

50. Mary Lilly and John Carlson. smellblind: a gene required for drosophila olfaction. Genetics, 124(2):293–302, 1990.

51. Mattias C Larsson, Ana I Domingos, Walton D Jones, M Eugenia Chiappe, Hubert Amrein, and Leslie B Vosshall. Or83b encodes a broadly expressed odorant receptor essential for drosophila olfaction. Neuron, 43(5):703–714, 2004.

52. RP Sharma. Light-dependent homosexual activity in males of a mutant of drosophila melanogaster. Experientia, 1977.

53. Peter R Michael, Danvers E Johnston, and Wilfrido Moreno. A conversion guide: Solar irradiance and lux illuminance. Journal of Measurements in Engineering, 8(4):153–166, 2020.

54. Shuying Gao, Shin-ya Takemura, Chun-Yuan Ting, Songling Huang, Zhiyuan Lu, Haojiang Luan, Jens Rister, Andreas S Thum, Meiluen Yang, Sung-Tae Hong, et al. The neural substrate of spectral preference in drosophila. Neuron, 60(2):328–342, 2008.

55. Inês MA Ribeiro, Michael Drews, Armin Bahl, Christian Machacek, Alexander Borst, and Barry J Dickson. Visual projection neurons mediating directed courtship in drosophila. Cell, 174(3):607–621, 2018.

56. A Frankel and G Brousseau. Drosophila medium that does not require dried yeast. Drosophila Information Service, 43:184, 1968.

57. Junko Kasuya, Atulya Iyengar, Hung-Lin Chen, Patrick Lansdon, Chun-Fang Wu, and Toshihiro Kitamoto. Milk-whey diet substantially suppresses seizure-like phenotypes of parashu, a drosophila voltage-gated sodium channel mutant. Journal of neurogenetics, 33(3):164–178, 2019.

58. Chun-fang Wu and Fulton Wong. Frequency characteristics in the visual system of drosophila. genetic dissection of electroretinogram components. The Journal of general physiology, 69(6):705, 1977.

59. Atulya Iyengar and Chun-Fang Wu. Flight and seizure motor patterns in drosophila mutants: simultaneous acoustic and electrophysiological recordings of wing beats and flight muscle activity. Journal of neurogenetics, 28(3-4):316–328, 2014.

