## Supplemental Files for "Intense light unleashes male-male courtship behavior in wild-type *Drosophila*"

**Supplemental Information: Ueda et al., 2023**

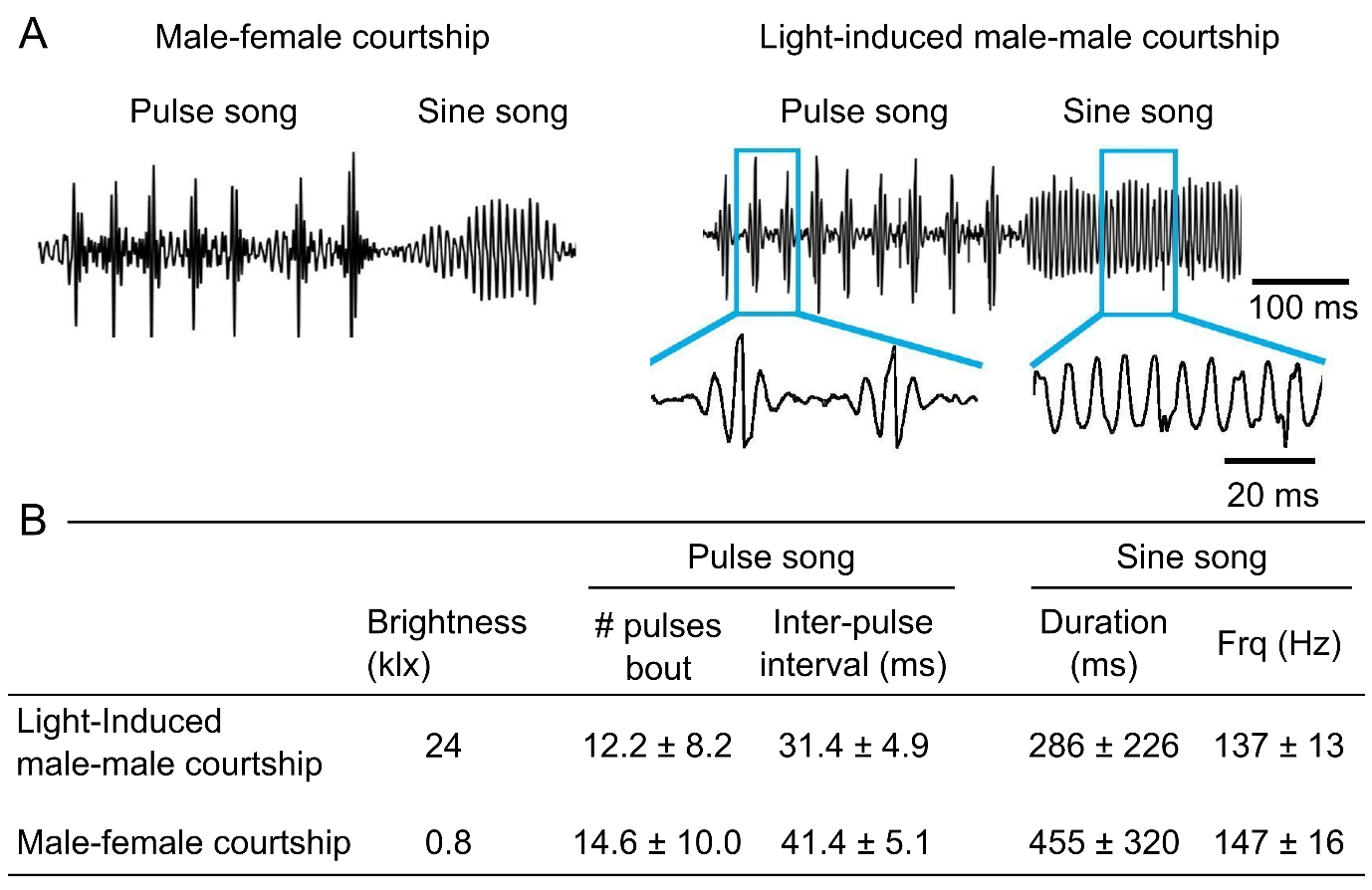

Figure S1. Comparisons between male-female courtship song and light-induced male-male courtship song. (A) Sample audio traces (see Methods). Intense light (24 klx) triggered pulse and sine songs in male-only arena (8 CS flies), resembling qualitatively songs recorded in mixed (CS males & females) arenas under low light (0.8 klx). (B) Parameters of courtship songs. Male-male courtship: 2 ROIs, 8 CS flies/ROI, 34-35 days old. Male-female courtship: CS, 3 ROIs, 3-8 males/ROI, 49 days old, and 2-4 virgin females/ROI, 5 days old. # samples: sine song = 11 for male-male and 10 for male-female courtship; pulse song = 68 for male-male and 18 for male-female courtship. Mean ± SD. Note overlaps between male-male and male-female courtship song parameters.

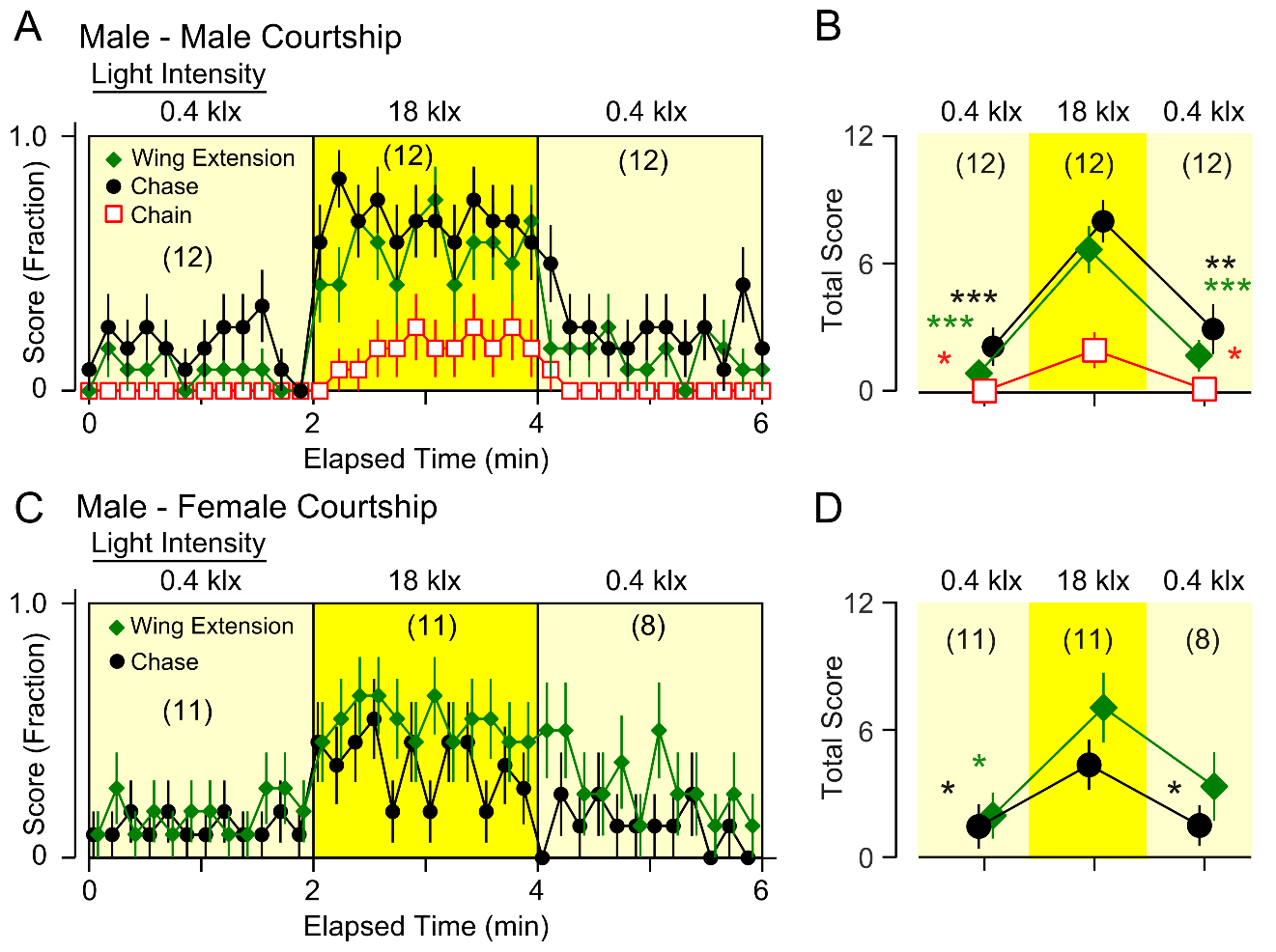

Figure S2. Intense-light effects on male-male and male-female courtship behaviors. (A, B) Kinematics and scores of light-induced male-male courtship (8 CS males/ROI), reproduced here from Figure 1D,E for comparison. (C) Kinematics of light-enhanced courtship behaviors between males and females. 4 CS males and 4 *w^1118^* females were used in each arena to facilitate identification of sex. Courtship behaviors between males and females was also increased by intense-light illumination (18 klx), but unlike male-male courtship, switching back to low light (0.4 klx) caused more gradual reduction in the male-female courtship scores. (D) Scores in C are summed over 2-min periods to produce the total score. Error bars indicate SEM. # ROIs are indicated in parenthesis. One-way ANOVA, Bonferroni-corrected t-test *post hoc* was applied for comparisons between 1^st^ low-light and high-light, as well as between high-light and the 2^nd^ low-light. **p <* 0.05, ***p <*0.01, ****p <*0.001.

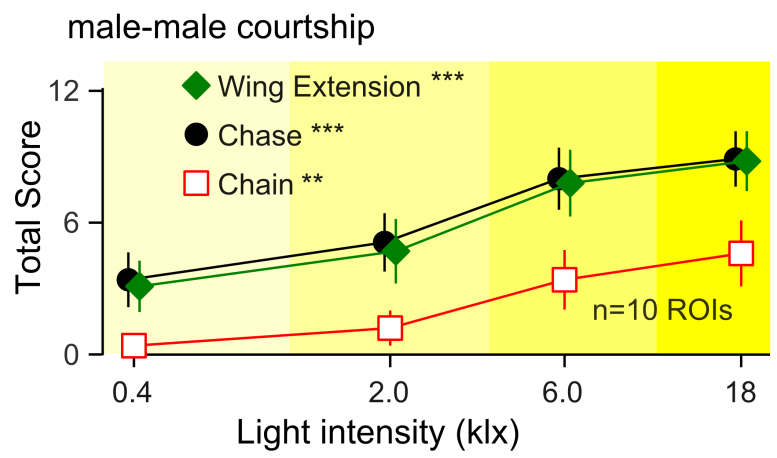

Figure S3. Light-intensity dependence of male-male courtship behavior among 8 male flies in an observation arena. Wing extension, chase, and chain were scored for 2 min under different illumination intensities of 0.4, 2.0, 6.0, and 18 klx. *** and ** indicates *p*< 0.001 and 0.01, respectively (Spearman’s Rank Test).

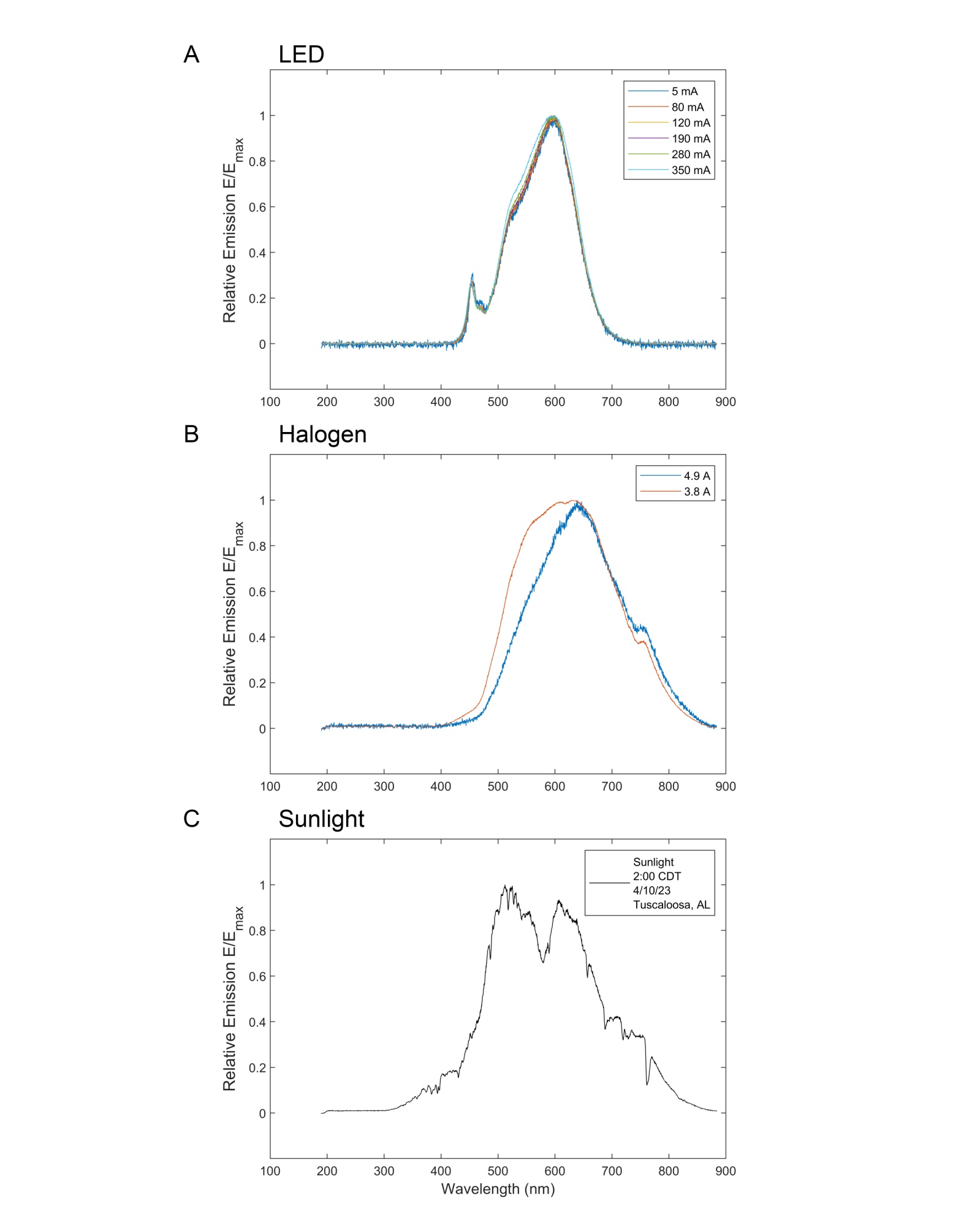

Figure S4 Emission spectra of light sources. (A) LEDs used in Figures 1-3 (see Methods for specifications) were driven by varying current intensities and the spectra across the 189 – 880 nm band was monitored. Data are normalized by the peak emission value (i.e. E(λ)/E_peak_ for wavelength λ). 8 mA corresponds with the ‘low’ intensity condition and 350 mA corresponds with the ‘high’ light intensity. (B) Data from the halogen incandescent bulb used in Figure 2C (see Methods). 3.8 A current corresponds with ‘low’ intensity and 4.9 A corresponds with ‘high’ intensity’. (C) Sunlight spectrum captured on a sunny day (2PM, April 10, 2023) in Tuscaloosa, AL, USA.

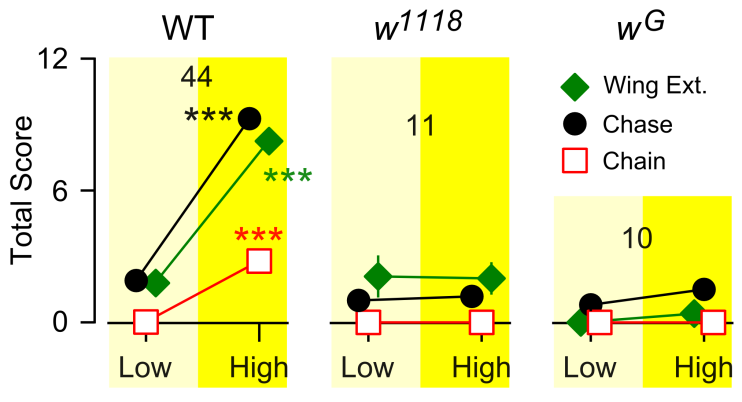

Figure S5. Lack of light-induced male-male courtship behavior in *w* mutants. 8 white-eyed male flies were introduced into the observation arena and their courtship behavior was recorded under low illumination (0.4 klx) for 2 min and then under high-intensity illumination (18 klx) for 2 min.*** *p*< 0.001, paired t-test (high vs. low). WT data replotted from Figure 2A for comparison.

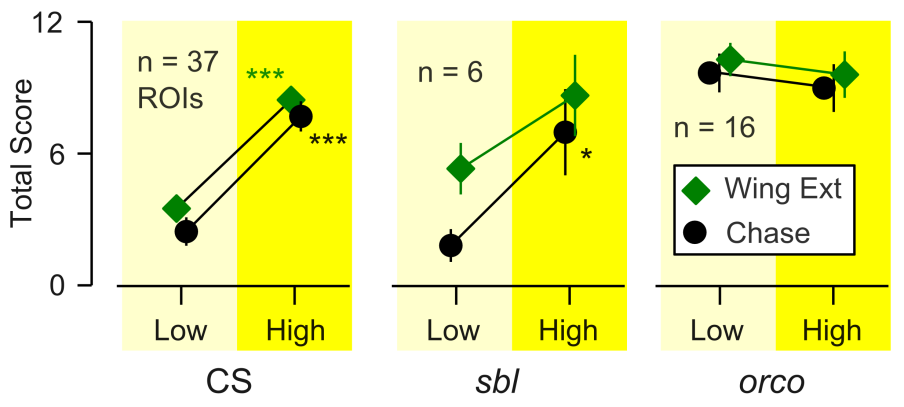

Figure S6. Effects of intense light on male-female courtship behaviors of *sbl* and *orco* mutant flies. CS data are reproduced from Figure 2B for comparison. Combined data from *sbl^1^* (n=5) and *sbl^2^* (n=1) as well as *orco^1^* (n = 6) and *orco^2^* flies (n = 10) are shown. Error bars indicate SEM. Low illumination (0.4 klx) for 2 min followed by high-intensity illumination (18 klx) for 2 min.

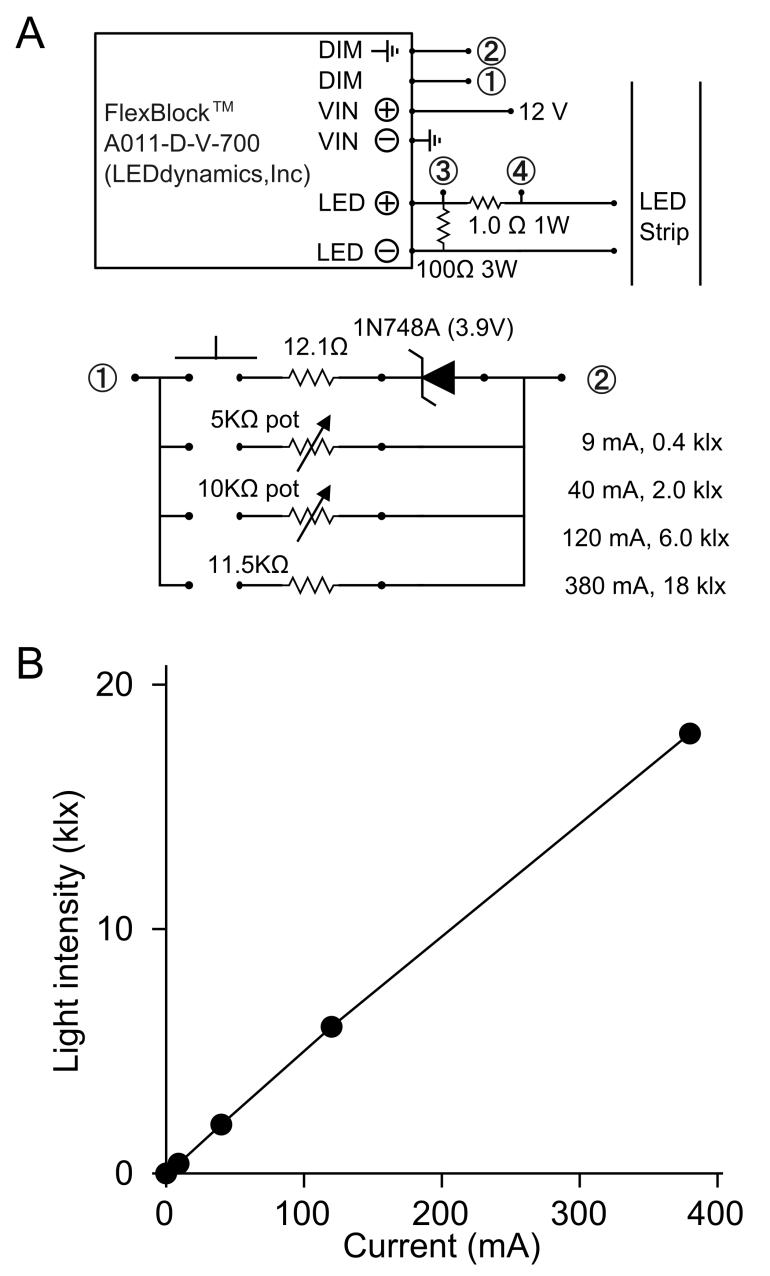

Figure S7. (A) Circuit diagram of the current regulator used to control the light output of the LED strip (STN-A30K80-A3A-10B5M-12V, SuperBrightLEDs.com) in the behavior chamber. The two potentiometers in A were used to obtain the intended middle levels of brightness (2.0 and 6.0, between 0.4 and 18 klx). (B) Current-brightness correlation in the behavior chamber. Current was measured as a voltage drop between the terminals 3 and 4 in A. The upper and lower control components in A are connected at terminals 1 and 2.

Table S1. Kinematic parameters of WT, *norpA*, and *orco* under low and intense light conditions.

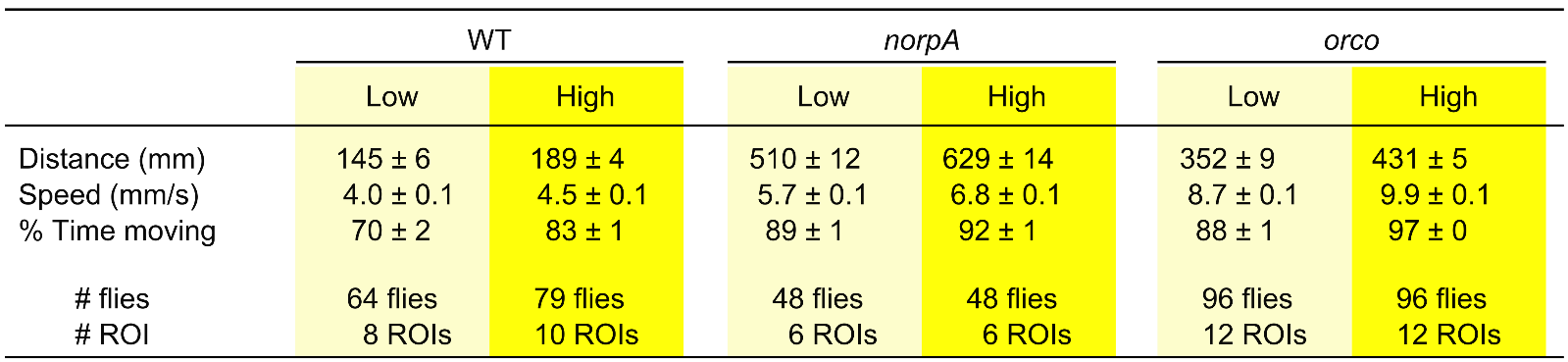

Notes: All genotypes showed statistically significant differences between low and high light conditions, one-way ANOVA, *post hoc* t-test (*p*<0.001). Also, these parameters in *norpA* and *orco* were different from WT under both low and high light conditions (*p*<0.001, Bonferroni-corrected t-test). SEM is shown.

Table S2. Male-male courtship behaviors of WT and *orco* mutant flies under white and blue light illumination.

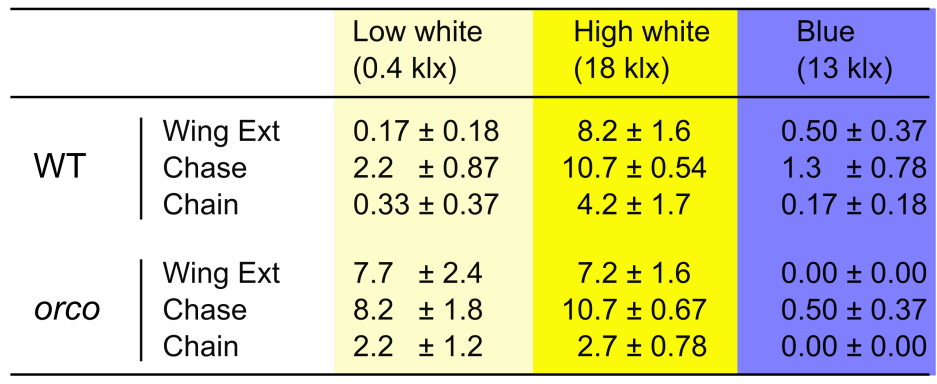

Notes: male-male courtship behaviors were suppressed under blue light (13 klx) following high white (18 klx), indicating that male-male courtship of *orco* is subjected to visual control and can be modified by visual input. # ROI: WT, n = 6; data from *orco^1^* (n = 3) and *orco^2^* (n = 3) are combined. Courtship behaviors were observed under 2-min low white, followed by 2-min high white, and then 2-min blue light illumination. Mean ± SEM. The criterion of ‘chain’ event was three or more animals in a row for this data set.

Table S3. Eye color and male-male courtship behaviors in *orco* homo- and hetero-zygotes and genetic interactions with visual mutations.

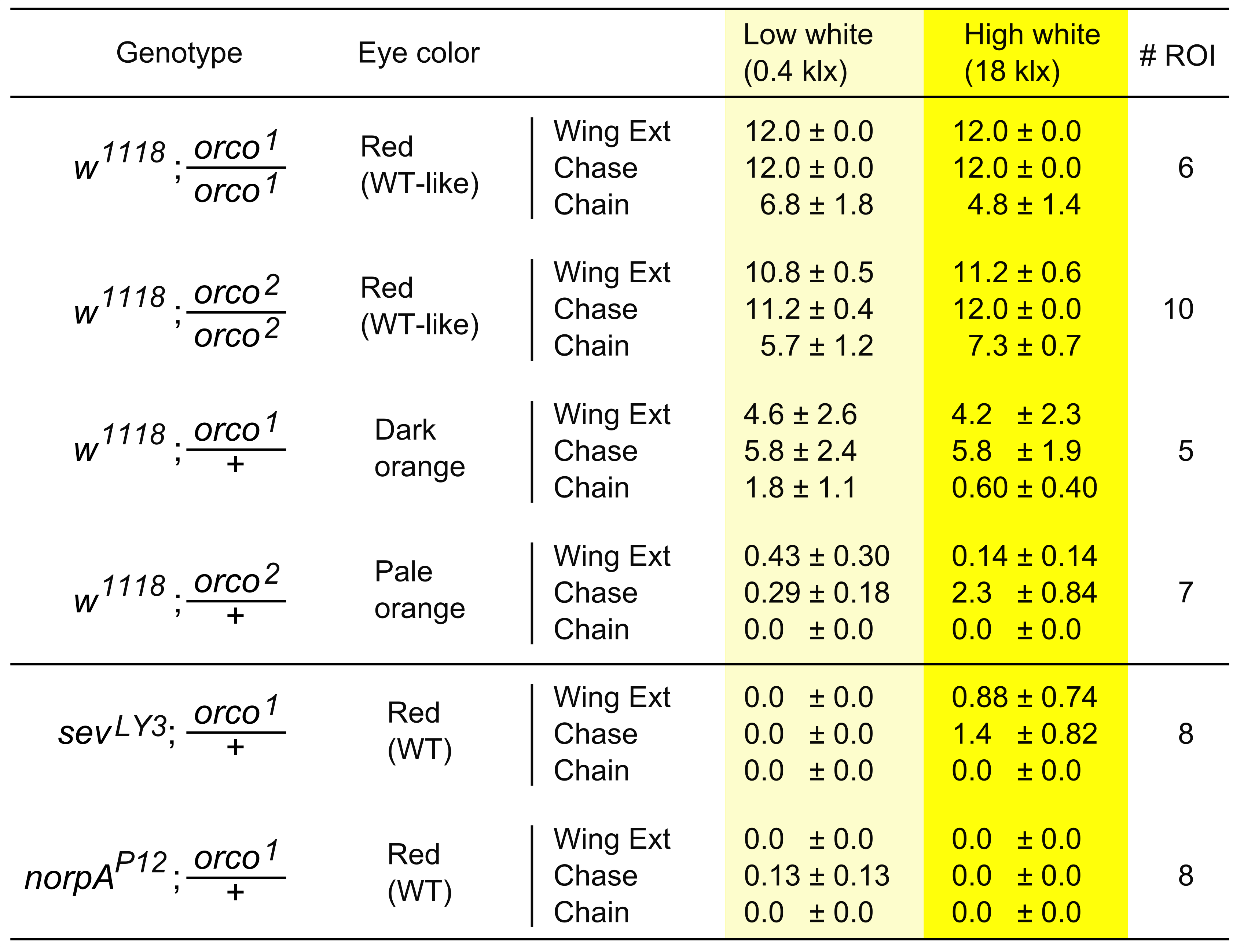

Notes: Higher behavioral scores appear to correlate with eye color. The number of copies of *mini-w^+^* (1 set in *orco*/+ heterozygotes and doubling in *orco*/*orco* homozygotes) determines the expression levels of eye-color pigments and thus the darkness of eye color. *w*; *orco*/*orco* shows darker eye color and higher behavioral scores than w; *orco*/+. (Behavioral scores of *orco* homozygotes are duplicate from Figure 2 for comparison.) The lower section of this table demonstrates a dominant role of visual processing in male-male courtship. When combined with visual mutations *sev* or *norpA*, the courtship behaviors of *orco^1^*/+ were greatly suppressed to the levels of *sev* and *norpA* (Figure 2), even though their eye color was indistinguishable from WT.

Table S4. Summary of statistical analyses in this paper.

| Figure | Test | Result |  | Deg. Freedom |
| --- | --- | --- | --- | --- |
| 1D | One-Way ANOVA (chasing) | F = 9.58 | p = 0.00052298 | 2/33 |
|  | *post hoc* paired t-test (chasing, low initial vs high) , holm-bonferroni corrected | p = 0.00004242 |  | 11 |
|  | *post hoc* paired t-test (chasing, low initial vs low final) , holm-bonferroni corrected | p = 0.096 |  | 11 |
|  | *post hoc* paired t-test (chasing, high vs low final) , holm-bonferroni corrected | p = 0.00004941 |  | 11 |
|  | One-Way ANOVA (wing ext) | F = 14.33 | p = 0.000033081 | 2/33 |
|  | *post hoc* paired t-test (wing ext, low initial vs high) , holm-bonferroni corrected | p = 0.00004291 |  | 11 |
|  | *post hoc* paired t-test (wing ext, low initial vs low final) , holm-bonferroni corrected | p = 0.241 |  | 11 |
|  | *post hoc* paired t-test (wing ext, high vs low final) , holm-bonferroni corrected | p = 0.00041522 |  | 11 |
|  | One-Way ANOVA (chaining) | F = 4.85 | p = 0.0142 | 2/33 |
|  | *post hoc* paired t-test (chaining, low initial vs high) , holm-bonferroni corrected | p = 0.045 |  | 11 |
|  | *post hoc* paired t-test (chaining, low initial vs low final) , holm-bonferroni corrected | p = 0.3388 |  | 11 |
|  | *post hoc* paired t-test (chaining, high vs low final) , holm-bonferroni corrected | p = 0.0398 |  | 11 |
| 2A, CS | paired t-test (chasing, low vs high) | p = 7.58E-18 |  | 42 |
|  | paired t-test (wing ext, low vs high) | p = 4.89E-15 |  | 42 |
|  | paired t-test (chaining, low vs high) | p = 1.974E-6 |  | 42 |
| 2A, Berlin | paired t-test (chasing, low vs high) | p = 0.0000771 |  | 4 |
|  | paired t-test (wing ext, low vs high) | p = 0.0218 |  | 4 |
|  | paired t-test (chaining, low vs high) | p = 0.203 |  | 4 |
| 2B | paired t-test (chasing, low vs high) | p = 1.307E-7 |  | 35 |
|  | paired t-test (wing ext, low vs high) | p = 1.969E-8 |  | 53 |
| 2C, Incand | paired t-test (chasing, low vs high) | p = 0.0096 |  | 4 |
|  | paired t-test (wing ext, low vs high) | p = 0.0201 |  | 4 |
|  | paired t-test (chaining, low vs high) | p = 0.2102 |  | 4 |
| 2C, Sun | paired t-test (chasing, low vs high) | p = 0.0038 |  | 2 |
|  | paired t-test (wing ext, low vs high) | p = 0.0012 |  | 2 |
|  | paired t-test (chaining, low vs high) | p = 0.2152 |  | 2 |
| 2D, norpA | paired t-test (chasing, low vs high) | p = 0.4933 |  | 9 |
|  | paired t-test (wing ext, low vs high) | p = 0.0379 |  | 9 |
|  | paired t-test (chaining, low vs high) | p = 1.000 |  | 9 |
| 2D, sev | paired t-test (chasing, low vs high) | p = 0.7609 |  | 10 |
|  | paired t-test (wing ext, low vs high) | p = 0.6332 |  | 10 |
|  | paired t-test (chaining, low vs high) | p = 1.000 |  | 10 |
| Table S4. *Continued* | | |  |  |
| Figure | **Test** | **Result** |  | **Deg. Freedom** |
| 2E, 1x | paired t-test (chasing, low vs high) | p = 0.00001731 |  | 7 |
|  | paired t-test (wing ext, low vs high) | p = 0.0014 |  | 7 |
|  | paired t-test (chaining, low vs high) | p = 0.0203 |  | 7 |
| 2E, 2x | paired t-test (chasing, low vs high) | p = 0.00004580 |  | 8 |
|  | paired t-test (chaining, low vs high) | p = 0.0390 |  | 8 |
| 2F, orco | paired t-test (chasing, low vs high) | p = 0.0561 |  | 14 |
|  | paired t-test (wing ext, low vs high) | p = 0.2997 |  | 14 |
|  | paired t-test (chaining, low vs high) | p = 0.8165 |  | 14 |
|  | unpaired t-test (chasing, orco low vs CS low) | p = 3.0405E-18 |  | 58 |
|  | unpaired t-test(wing ext, orco low vs CS low) | p = 2.6552E-18 |  | 58 |
|  | unpaired t-test(chasing, orco low vs CS low) | p = 5.9018E-18 |  | 58 |
| 2F, sbl | paired t-test (chasing, low vs high) | p = 0.0105 |  | 5 |
|  | paired t-test (wing ext, low vs high) | p = 0.0147 |  | 5 |
|  | paired t-test (chaining, low vs high) | p = 0.0311 |  | 5 |
| Figure S2D | One-Way ANOVA (chasing) | F = 5.29 | p = 0.0138 | 2/28 |
|  | *post hoc* paired t-test (chasing, low vs high) holm-bonferroni corrected | p = 0.0148 |  | 6 |
|  | *post hoc* paired t-test (chasing, low vs low final) holm-bonferroni corrected | p = 0.05097 |  | 6 |
|  | *post hoc* paired t-test (chasing,high vs low final) holm-bonferroni corrected | p = 0.0254 |  | 6 |
|  | One-Way ANOVA (wing ext) | F = 4.16 | p = 0.0277 | 2/28 |
|  | *post hoc* paired t-test (wing ext, low vs high) holm-bonferroni corrected | p = 0.0144 |  | 6 |
|  | *post hoc* paired t-test (wing ext, low vs low final) holm-bonferroni corrected | p = 0.1900 |  | 6 |
|  | *post hoc* paired t-test (wing ext, high vs low final) holm-bonferroni corrected | p = 0.1136 |  | 6 |
| Figure S3 | Spearman's Rank (wing ext) | p = 0.00011 |  | 38 |
|  | Spearman's Rank (chasing) | p = 0.00012 |  | 38 |
|  | Spearman's Rank (chasing) | p = 0.0063 |  | 38 |
| Figure S5 | paired t-test (w1118 chasing, low vs high) | p = 0.3419 |  | 10 |
|  | paired t-test (w1118 wing ext, low vs high) | p = 0.7496 |  | 10 |
|  | paired t-test (w1118 chaining, low vs high) | p = 1.000 |  | 10 |
|  | paired t-test (wG chasing, low vs high) | p = 0.2639 |  | 9 |
|  | paired t-test (wG wing ext, low vs high) | p = 0.0592 |  | 9 |
|  | paired t-test (wG chaining, low vs high) | p = 1.000 |  | 9 |
| Figure S6 | paired t-test (orco, chasing low vs high) | p = 0.6273 |  | 4 |
|  | paired t-test (orco, wing ext low vs high) | p = 0.6024 |  | 4 |
|  | paired t-test (sbl, chasing low vs high) | p = 0.0339 |  | 14 |
|  | paired t-test (sbl, wing ext low vs high) | p = 0.1600 |  | 14 |
